# Stochasticity explains non-genetic inheritance of lifespan and apparent trade-offs between reproduction and ageing

**DOI:** 10.1101/2023.03.06.531380

**Authors:** Elizabeth D Drake, Mirre J P Simons

## Abstract

Stochastic effects are central to the biology and demography of ageing. Genetically identical individuals do not all die at the exact same time but show a distribution of lifespan. Although such effects are appreciated, any cascading effects from stochastic effects of ageing are underappreciated. We show here that genetically identical female flies (*Drosophila melanogaster*) that live long, produce longer lived daughters. In line with previous work, we also find that daughters born to older mothers are shorter lived, also termed the Lansing effect. We further show that longer-lived flies produce less offspring, suggesting an apparent trade-off due to stochastic effects alone. We explain these effects using an extension of the reliability theory of ageing by dichotomising ageing physiology in reproduction and lifespan supporting units. These simple models reproduce non-genetic inheritance of lifespan, the Lansing effect and trade-offs between reproduction and lifespan. Our work implies that if non-genetic inheritance of lifespan is widespread it explains the generally low heritability of this trait. Furthermore, trade-offs between performance, e.g. reproduction, and lifespan may be less widespread than predicted by evolutionary biology of ageing, stemming from stochasticity rather than differential investment. Anti-ageing treatments therefore come without any unintended costs to other physiology, a perceived risk that limits translation of these treatments to humans.

## Introduction

Ageing is existential, but poorly understood in both its evolutionary and mechanistic biology^1–4^. In a population, not all individuals appear to age at the same rate^5^ in terms of lifespan and reproduction^6^. Moreover, it is not always evident that these two traits are negatively correlated, as some individuals appear to both show high reproduction and longevity in a population, often termed quality effects^7–9^. Understanding reproduction and lifespan is critical to understand ageing at a fundamental level as they are most closely related to Darwinian fitness. When we understand such variation we can start to understand how selection acts on ageing and partition environmental from genetic effects. Surprisingly, however, for both reproductive output and lifespan a large amount of variation has been observed in individuals of the exact same genotype kept in the exact same environment^10–12^. In the absence of genetic and environmental effects, stochastic and parental effects remain.

Stochastic effects in ageing are evident, as the resultant phenotype lifespan is inherently variable. Lifespan follows from a presumed genetically determined mortality risk function with age^13,14^. How reproduction changes with age, often termed reproductive senescence, has also been shown to vary amongst individuals in human and other animal populations^15,16^. In addition to this there are mechanistically unexplained effects of parental age on offspring lifespan and reproduction^17,18^. Non-genetic inheritance of how organisms age is thus documented but not understood. Here we show non-genetic inheritance of lifespan using an inbred population of fruit flies (*Drosophila melanogaster*). Mothers that lived the longest produced long-lived daughters.

Differences in Darwinian fitness traits are often explained as differences in quality^7,19^. The term quality encompasses differences in local adaptation or long-lasting effects of early environmental effects. In line with this, in many animal populations, individuals with high reproductive rates or high lifetime reproductive success are also often the most long-lived ^6,7,20^. We find, however, that mothers that are long-lived are not typical high quality individuals, as longevity was associated with reduced reproductive output. Across genetically identical individuals we thus find evidence of an apparent trade-off between reproduction and lifespan. Such trade-offs have been fundamental to explaining life-history theory and the evolution of ageing^1,21,22^. Here we provide a novel explanation to non-genetic inheritance of lifespan and parental age effects, and apparent trade-offs between reproduction and lifespan, using the reliability theory of ageing.

### Theoretical Model

Ageing can be modelled in various ways. The earliest models described ageing as an exponential mortality risk curve^13^ and stochastic effects have been central in models of ageing, especially those describing mortality^23^. Mortality of many species can be accurately described using a simple two parameter exponential risk model such as the Gompertz equation^24,25^. Perhaps more importantly treatment effects^24,26,27^ and differences between human populations can be similarly captured by these models^28^. Risk, and thus stochasticity within the population, is fundamental to how we understand ageing. In the underlying biology of ageing we also find these ideas, with damage accumulation with age as central^29^ to all physiology implicated in ageing. Although the central role of damage accumulation with age has been challenged by quasi-program and developmental explanations of ageing^30^, these theories currently fail to explain why exponential increasing risk with age appears central to ageing.

A theory that has been underappreciated but provides an excellent quantitative embedding of ageing related biology, is the reliability theory of ageing. This theory states that organisms consist of units that fail at a set pace (termed ‘failure rate’), and that organisms have a level of redundancy built in (termed ‘units’)^31,32^, and when all redundancy is lost the system fails. Importantly, this theory explains why mortality risk plateaus, and converges at old age^31^.

More recent applications of this theory have explained why biomarkers of physiological system redundancy reduce in explanatory power with age^33^. Here we use the reliability theory of ageing to explain the non-genetic inheritance and maternal age effects we find in the fruit fly. In addition, our extension of the reliability theory of ageing predicts apparent trade-offs between physiological domains of ageing, that result from stochasticity rather than differential investment. The latter is assumed in the disposable soma theory and life-history theory, although evidence of such investment is lacking^2,34^.

We assume that physiology consists of units that fail with age as in the original reliability theory of ageing. Of these units of physiology we now assume some support physiology that prevents mortality and others support reproductive output. Such compartmentalization in physiology separating aspects that cause mortality, from others that decline with age are probably prevalent. Ageing of different aspects of physiology is expected to be related to mortality risk in varying degrees. For example, we believe wrinkles to not be causally linked to mortality even though the presence of wrinkles is a correlate of age and thus mortality risk. Other physiology, such as immune functioning might be more of a central role in overall physiology and might thus determine, for example, age-related reproductive performance or endurance, as well as age-related mortality risk. Of interest to us here is how these physiological domains are expected to age in relation to each other in a genetically identical population.

In the original models of ageing using reliability theory each unit of physiology has an independent chance of failing over time. Thus without any variation in failure rate of the system’s components and of how much redundancy the system has, each physiological domain will age independently as each failure is an independent chance event. The alternative, not explored in these models before, is one of constant damage to the system but with the damage falling on components of the system in a stochastic fashion. This generates an inverse relationship between physiological domains as damage falling on one part of physiology will not fall on the other. When such damage is attributed to the system in a draw with replacement, i.e. a failed unit that is damaged can absorb damage, this model behaves similarly to original reliability models of ageing. Mortality converges to the risk of the rate of failure divided by all redundancy in the system.

Reliability theory models incorporating compartmentalisation of physiology and constant damage applied through a draw with replacement at each timepoint were simulated using R. Our model is intended to aid interpretation of our findings rather than be parameterized to observations. We modelled a system of 10 physiological units, with 5 sustaining reproduction, and 5 sustaining lifespan (Figure 1A) in a population comprising 10,000 individuals with damage of one unit per time (redraw across the 10 units). The level of units left in reproduction we propose is a metric of the ability to sustain reproduction. When all units are damaged in the lifespan sustaining system, an individual dies (Figure 1D). The mutual exclusivity of damage distribution on this physiological network with stochastically allocating a constant level of damage on the network, means some individuals in the population will age disproportionally in either physiological domain (Figure 1B & 1C). Individuals that lose system integrity on the reproduction side more rapidly, have a larger proportion of their lifespan sustaining physiology left intact, and *vice versa* (Figure 1B & Figure 1C). Stochasticity thus leads to an apparent trade-off between reproductive output and lifespan across individuals that are intrinsically physiologically identical (Figure 1E, F).

**Figure 1.**
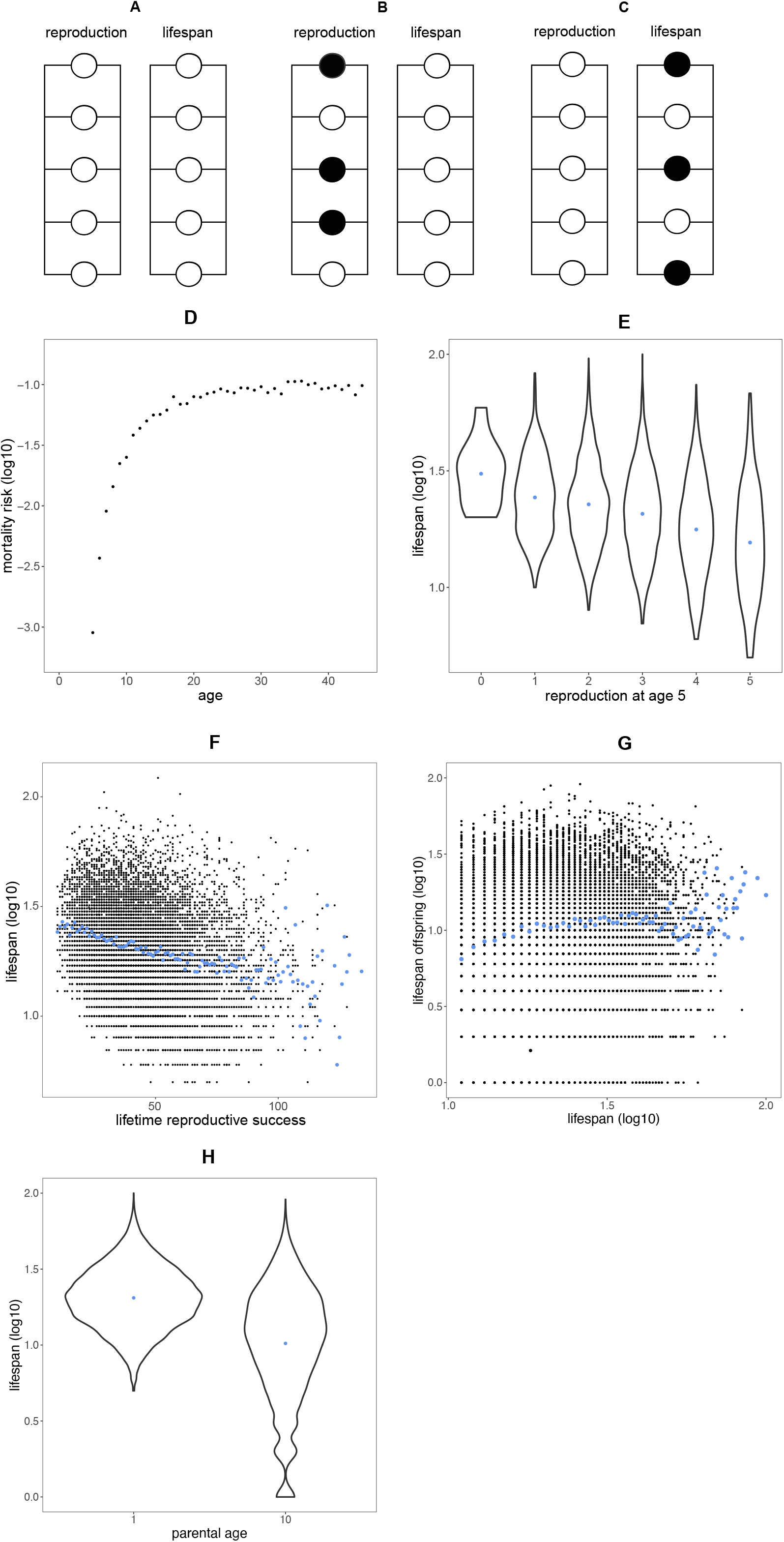
**A**. Reliability theory model of ageing partitioning physiology between reproduction and longevity. Damage is drawn as one unit failing per time across all ten units (including those that have already been damaged). Some individuals simply due to stochasticity age disproportionately either in reproduction (**B**) or lifespan sustaining physiology (**C**). The model behaves as a standard reliability theory of ageing model, with mortality increases semi-exponentially with age with mortality converging at the oldest ages (**D**). The partitioning between reproduction and lifespan but their shared susceptibility of one unit being hit by damage per time generates a negative relationship between reproduction (here number of units left equals reproductive output) and lifespan (**E**). This effect is reminiscent of the reproduction, lifespan trade-off central to evolutionary biology and mechanisms of ageing, but notably only came about through stochastic effects. Moreover this relationship extends into total lifetime reproductive success with individuals with long lifespans producing less offspring in total. When non-genetic inheritance is incorporated into this model by assuming lifespan sustaining physiology is inherited somehow additional insight is gained. This simple stochastic model generates a positive relationship between lifespan of the parent and lifespan of the offspring (**G)** and a negative relationship between parental age and offspring lifespan (**H**).

We can further apply this model to understand the effects of parental age on offspring fitness^35^ and lifespan^17^, also termed the Lansing effect. We can hypothesise either part of physiology, the reproduction or lifespan sustaining component is passed down through non-genetic inheritance. When organisms would pass on their age-dependent level of life-sustaining redundancy this generates a relationship between parental age and offspring lifespan, because redundancy is lost as a function with age. Furthermore, such a hypothesised non-genetic inheritance mechanism would generate a positive relationship between lifespan of the parent and the offspring. As the level of redundancy left at a given parental age is a predictor of both parental and offspring survival. These assumptions show that similar physiology could underlie both parental age effects and parental offspring lifespan relationships.

## Results & Discussion

Mothers that lived long produced long-lived daughters (*r*_s_ = 0.49, P = 0.006, Figure 2A, 2D, HR(coxme) = −0.016± 0.0059, P = 0.005 per day of maternal lifespan). Daughters born to mothers at older ages showed reduced longevity (HR(coxme) = 0.11± 0.02, P < 0.001 per day of maternal age, Figure 2B, 2E). Mothers that lived long had less reproductive output (r_s_ = 0.49, P = 0.028, as measured across both ages, Figure 2C). Note that any of these effects are unlikely due to population size differences in the growing vials, as total pupal case numbers per vial did not correlate to the resulting lifespan of offspring from those vials (r_s_ = 0.048, P = 0.74).

**Figure 2.**
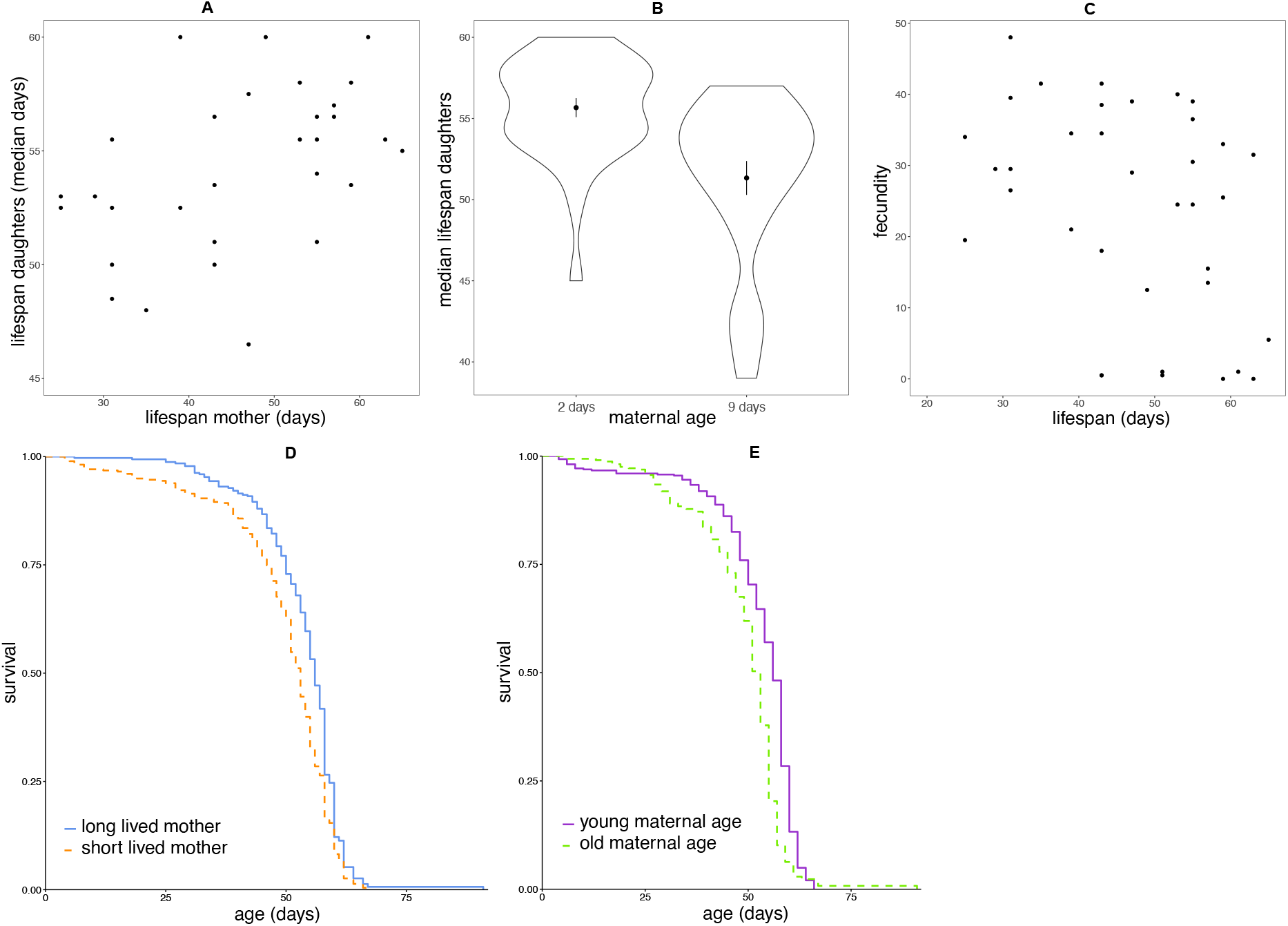
Non-genetic inheritance of longevity in inbred fruit flies. **A, D**. Mothers that were long lived produced daughters that were long lived (median). **B, E**. Daughters born to mothers of older ages survived less long. **C**. Mothers that lived long produced less offspring.

The fly experiments indicate that in a highly inbred genetic line, stochastic effects inherent to the ageing process can explain several findings that currently lack a mechanistic explanation. Offspring from older mothers show truncated lifespans, also termed the Lansing effect^17,36^. Similarly, parental and offspring lifespan correlations are often interpreted in a genetic context only^37^. Our findings suggest that part of the correlation between parent and offspring lifespan could originate from the same physiology that underlies the Lansing effect. Note, that such parental effects would affect twin studies^38^ as well as parent-offspring correlation and pedigree-based approaches to heritability of lifespan. When non-genetic inheritance of lifespan is ubiquitous and due not to what is usually interpreted as environmental effects, but to stochastic effects alone, the realised heritability of a trait and thus its response to selection reduces^39^.

A final important observation from our work is that trade-offs can emerge from stochastic effects during ageing alone. Individuals that age fast in one physiological domain supporting e.g. lifespan, age less fast in another physiological domain, e.g. reproduction. Trade-offs between reproduction and longevity are central to life history but when tested experimentally lack empirical support, especially in naturalistic settings^7,40^. As an explanation for this, differences in phenotypic quality, allowing some individuals to show both high reproductive output as well as longevity, has been suggested to mask trade-offs and hence limit selection^40,41^. In contrast, we show here that apparent trade-offs between traits can emerge from stochastic ageing in inbred flies and in our theoretical models. Thus, in both directions of the fitness landscape natural selection is limited in its potential to select for sharp trade-offs. The assumed strength of the trade-off between reproduction and longevity, central to the biology of ageing, could thus be overstated. We should therefore not be forced to assume costs of anti-ageing treatments to other physiological domains, although dictated by life history doctrine. This idea fits with observations of cost-free longevity extension^42^. When costs of longevity treatments are not presumed, translating findings from biology of ageing research to the clinic increases in feasibility^43,44^.

## Methods

### Experiments in the fly

Fruit flies of the standard inbred laboratory (*yw*) stock were used for the experiments^45^. All experiments were conducted on our standard lab diet (‘rich’), 8% yeast^46^. Mothers were grown together in one bottle to ensure shared environmental effects and these were mated together for two days after eclosion. Mothers (n=40) were then single housed in vials until they died to record their lifespan (3 flies were lost to follow up). Food vials were changed every two days when a census was taken. Daughters of these mothers were collected from vials when mothers were 2 to 3 days old and 9 and 11 days old, were mated in a vial for 2 days (presence of males was confirmed), and then recorded as a population for longevity using demography cages (n=1 to n=31 per cage, maximum of one cage per collection point). Only intact offspring were used and individuals were censored if they were stuck to the food or escaped from the cage during handling^2^. Total offspring produced from the individual mothers was recorded by counting the pupal cases at both collection timepoints. Data was analysed using spearman rank correlations, linear-mixed effects models and cox proportional hazard models that included random terms for cage and mother and right-hand censoring. Results from the spearman rank correlations and cox models are presented, as the linear mixed effect models gave qualitatively similar results.

## Author contributions

EDD designed and conducted the experiments and reviewed the manuscript. MJPS designed the experiments, produced the model and wrote the first draft of the manuscript.

## Acknowledgements

MJPS is supported by a Sir Henry Dale Fellowship (Wellcome and Royal Society; 216405/Z/19/Z) and an Academy of Medical Sciences Springboard Award (the Wellcome Trust, the Government Department of Business, Energy and Industrial Strategy (BEIS), the British Heart Foundation and Diabetes UK; SBF004\1085). For the purpose of Open Access, the author has applied a CC BY public copyright licence to any Author Accepted Manuscript version arising from this submission. We thank Laura Hartshorne and Gracie Adams for technical support. We thank Marc Tatar for supplying the fly stock used.

